# A Scalable fMRI Estimate of Basal Ganglia Brain Tissue Iron for Use in Developmental and Translational Neuroscience

**DOI:** 10.64898/2026.04.10.717850

**Authors:** Holly Sullivan-Toole, Ashley C. Parr, Carina Heller, Brenden Tervo-Clemmens, rae McCollum, Amar Ojha, Eric Feczko, Erik Lee, Will Foran, Finnegan J. Calabro, Beatriz Luna, Bart Larsen

## Abstract

Dopaminergic (DA) function and basal ganglia neurobiology are central to reward learning, motivation, and cognitive control, and dysregulation of these systems contributes to neuropsychiatric conditions that emerge during development. Adolescence is marked by profound reorganization of DAergic basal ganglia circuitry, yet direct *in vivo* assessment of the DA system remains limited in youth. Brain tissue iron is a developmentally sensitive marker of DA-related neurobiology that can be measured non-invasively via magnetic resonance imaging (MRI). Iron is an essential co-factor for DA synthesis and a foundational metabolic resource that supports cellular metabolism, myelination, and energetic demands of the basal ganglia. T2*-weighted echo-planar imaging (EPI), collected during functional MRI (fMRI), is sensitive to magnetic susceptibility of non-heme brain iron. Leveraging this property, we demonstrate the validity and broad applicability of an iron-sensitive metric that can be derived from conventional single-echo fMRI: ΔR2*. In a longitudinal developmental dataset (N = 151; age range 12–31), ΔR2* showed high reliability, strong longitudinal stability, and validity via robust convergence with established quantitative relaxometry-based iron measures (R2* and R2’). Critically, ΔR2* can be retrospectively estimated from extant fMRI data and derived in large-scale consortium data repositories, demonstrated here in the Adolescent Brain and Cognitive Development (ABCD) baseline cohort (N = 8,366; ages 9–11). We show that ΔR2* captures known age-related increases in basal ganglia iron, highlighting neurodevelopmental sensitivity at population-scale. Together, these findings establish ΔR2* as a reliable, widely accessible marker of basal ganglia iron, enabling scalable investigation of lifespan trajectories and neuropsychiatric risk in existing and future datasets.

## INTRODUCTION

Adolescence is a critical period of neurodevelopment characterized by substantial remodeling of neural systems that support reward learning, motivation, and cognitive control^1,2^. Studies in humans and animals demonstrate that dopamine (DA)-dependent processes within the basal ganglia undergo pronounced developmental reorganization during this period, with functional consequences for cognitive and affective behaviors, including adolescent peaks in sensation seeking and risk-taking behaviors across species and cultures^3^. Dysregulation of basal ganglia DA processes contributes to neuropsychiatric conditions that frequently emerge during adolescence, including substance use^4,5^ and psychosis^6^. In animal models, increasingly sophisticated molecular-, circuit-, and systems-level approaches reveal dynamic shifts in basal ganglia DA physiology throughout adolescence – including changes in receptor expression, presynaptic availability, synthesis capacity, and phasic firing patterns^3,7–12^. Together, these findings highlight DA-related neurodevelopment as a central mechanism underlying adolescent behavioral phenotypes and psychiatric vulnerability.

However, direct assessment of DA and basal ganglia neurobiological development in youth remains challenging. Positron Emission Tomography (PET) offers unparalleled *in vivo* estimates of DA receptor availability, synthesis capacity, and presynaptic function, but is largely infeasible in healthy pediatric populations due to its invasive nature (use of intravenous injection) and ethical considerations surrounding radiation exposure^13^. Brain tissue iron has emerged as a compelling, non-invasive marker of DA-related neurobiology that also captures broader aspects of neural metabolic capacity. Tissue iron (ferritin) is heterogeneously distributed throughout the brain but is most highly concentrated in the DAergic regions including the basal ganglia and midbrain nuclei^14–17^, which reflects tissue iron’s essential roles in DA neurophysiology. Tissue iron serves as an essential co-factor for tyrosine hydroxylase^18,19^ – the rate-limiting enzyme in DA synthesis – and is enriched in mesostriatal DA-producing nuclei^19,20^. Iron is stored in oligodendrocytes, neurons, and microglia^21,22^, and is necessary for a myriad of enzymatic and metabolic functions essential to brain development – including mitochondrial and oxidative metabolism^23^, myelination^24,25^, and monoamine synthesis^18,19^. These processes collectively support the energetic demands of circuit maturation, synaptic signaling, and plasticity^26,27^ during development as well as the maintenance of neural integrity across the lifespan^21^. Tissue iron also influences glial function, neuroimmune signaling, and oxidative balance – processes increasingly recognized as central to developmental plasticity^26,27^ and long-term circuit health^21^. Thus, iron can be conceptualized as a foundational metabolic resource constraining both DA signaling and the broader capacity of basal ganglia circuits to generate and sustain functional responses. Brain tissue iron may thus play a significant role in DAergic models of neurodevelopment and neuropsychiatry, where insufficient or dysregulated iron availability may undermine DA synthesis and neural and circuit efficiency, contributing to the functional dysregulation observed in neurodevelopmental, neuropsychiatric, and neurodegenerative disorders.

Brain iron rapidly increases during childhood and adolescence, followed by slower accumulation through adulthood and aging^28–38^. These trajectories parallel major neurodevelopmental changes, including myelination^39,40^, synaptic refinement^41,42^, and the maturation of cortico-subcortical circuitry^43–45^, as well as adolescent reorganization of DA-sensitive circuits^43,46^. Iron deficiency during development has lasting effects on cognition and attention^47–49^, while excessive or dysregulated iron accumulation contributes to oxidative stress and neuroinflammation implicated in neurodegenerative disease, including Parkinson’s disorder^19,50^ and restless leg syndrome^51^. Altered basal ganglia brain iron levels have been reported across a range of neuropsychiatric and neurodevelopmental conditions, including ADHD^52,53^, mood disorders^54,55^, psychosis risk^56^, and substance use^35,57^. Collectively, these findings position iron homeostasis as a key determinant of healthy neurodevelopment and psychiatric vulnerability.

Because brain iron is paramagnetic, it influences the effective transverse relaxation (T2*) by impacting magnetic susceptibility and through its effect on local magnetic field inhomogeneity. This property allows brain iron to be quantified *in vivo* using specialized MRI acquisitions such as R2* and R2’, which index T2* relaxometry, or the rate of decay of the T2*-weighted signal. These relaxometry-based measures correspond closely to postmortem iron assays^25,58^, and work from our group has demonstrated both cross-sectional and longitudinal correspondence between estimates of brain iron assessed via quantitative relaxometry (R2’) and presynaptic vesicular DA storage measured using PET [^11^C]dihydrotetrabenazine (DTBZ)^38,59^. Developmental neuroimaging studies consistently report increases in MR-based indices of tissue iron accrual throughout the first two decades of life, with the rate of iron increase decelerating into adulthood^28–32^, paralleling DA-related developmental patterns observed in animal models^60^. Iron-sensitive MRI measures have further been linked to key adolescent neurobehavioral and neurocognitive processes, including reward sensitivity^33^, cognitive control^30,34,47^, and functional changes in mesocorticolimbic networks that support developmental declines in risk-taking from adolescence to adulthood^43^. Together, these findings support iron-sensitive MRI as a biologically meaningful, developmentally-sensitive, and non-invasive index of basal ganglia iron that reflects important aspects of DA neurobiology.

Despite the strong theoretical rationale for using MR-based indices of brain iron to index DA-related neurobiology in the basal ganglia, quantitative relaxometry (e.g., R2’, R2*) and susceptibility imaging require specialized multi-echo or susceptibility-weighted acquisitions that are not routinely collected or are not feasible in most developmental or clinical neuroimaging studies. Consequently, standard markers of tissue iron content cannot presently be directly quantified across many existing datasets – including large-scale cohorts such as the Adolescent Brain and Cognitive Development (ABCD) Study^61^ – limiting opportunities to model iron-related neurodevelopmental processes in large populations or to investigate iron as a mechanistic contributor to psychiatric risk at population scale.

fMRI, one of the most widely collected modalities in pediatric and clinical research, presents a scalable alternative. Relaxometry-based approaches generally collect sequences of T2*-weighted (T2*w) images at different echo times to quantify the rate of T2* decay over time (see^62^ for review), with the image contrast within each T2*w image reflecting the relative decay across the brain at a given echo time (TE). Iron shortens the T2* signal, increasing the rate of transverse relaxation through its effects on magnetic susceptibility and local field inhomogeneity, and therefore, regions with higher iron content will exhibit faster T2* relaxation relative to areas with less iron content^63–65^. The signal amplitude at a single TE therefore reflects T2* decay that has occurred by that time, such that single-echo signal intensity provides a proxy for the underlying iron-dependent T2* decay rate. Consequently, single-echo echo-planar imaging (EPI) acquisitions, which are common in standard fMRI approaches, yield T2*w images that are sensitive to regional variation in iron-sensitive T2* decay. While fMRI analyses typically focus on temporal fluctuations in the T2* signal over time (i.e., BOLD), the signal contrast of a single T2*w image can be used to derive a marker of relative local iron concentration levels in a given brain area^66^. This approach relies upon the comparison of the T2*w signal to a reference tissue (e.g., a whole-brain mask), generating a relative signal intensity, and builds upon prior work, including work from our group that has used a number of variations of T2*w contrast (e.g., time-averaged and normalized T2*w imaging (nT2*w)) as an iron-related marker in the brain^4,36,53^ and other organs (e.g., the liver^67^). Although the T2*w signal has been shown to correspond to brain iron content^68^ and DAergic neuron concentration in midbrain nuclei^69^, a systematic validation of these measures – including assessment of reliability, longitudinal stability, and convergence with established and validated quantitative relaxometry-based iron measures – has not been performed, limiting the broader adoption and translational utility of an fMRI-derived iron measure.

Here, we present and validate ΔR2* as a scalable, fMRI-derived estimate of basal ganglia tissue iron for existing developmental and translational neuroscience datasets. ΔR2* builds upon our prior investigations with the nT2*w contrast by applying a negative log transformation normalization— informed by recent work from Garzón and colleagues—that reflects the *difference* in R2* between a given tissue and the reference tissue and ensures that the metric scales positively with tissue iron concentration. We first establish construct validity by assessing the correspondence of ΔR2* with well-established quantitative relaxometry metrics. We then evaluate its reliability and longitudinal stability across both short-term (within-session) and long-term (multi-year) intervals. Next, we evaluate harmonization procedures enabling ΔR2* to be compared across sites, scanners, and longitudinal time points (e.g., months, years), addressing a major obstacle for large-scale, multi-site neuroimaging research. Finally, we test sensitivity to established brain iron developmental trajectories and to interindividual differences by characterizing age-related changes and sex differences in basal ganglia ΔR2* in the ABCD cohort – where dedicated iron neuroimaging is unavailable. To facilitate broad adoption, we provide a lightweight, open-source tool for deriving ΔR2* that is compatible with data processed with *fMRIPrep*, enabling researchers to prospectively and retrospectively investigate iron-related neurodevelopment and neuropsychiatric risk at scale.

## METHODS

### Processing pipeline evaluation and assessment of the validity and reliability of ΔR2*

We evaluated a series of processing pipelines designed to produce brain iron-related estimates derived from the T2*w signal contained in standard single-echo EPI fMRI data. Each step of the processing pipeline was exhaustively compared to evaluate the impact on the derived metrics. After arriving at a preferred processing pipeline, we further conducted a series of analyses to establish validity via correspondence with quantitative relaxometry-based brain iron measures and to establish reliability across repeated measurements spanning hours to years. All analyses related to pipeline evaluation, validity, and reliability were performed in an imaging dataset collected by the study team (see Sample 1 below), which has been reported on previously^31,33,38,43,70^. Sample 1 data and analysis code is available at https://github.com/hollysully/dR2star_validation/tree/main

### Sample 1 participants

Adolescent and young adult participants participated in a longitudinal MRI study (Luna - MH080243) that included acquisition of multiple runs of resting-state fMRI as well as quantitative relaxometry MRI. The study consisted of three longitudinal visits spaced approximately 18 months apart. At each visit, participants completed structural T1 imaging, relaxometry-based R2 and R2* acquisition, and two runs of resting-state fMRI. Restricting the study sample to those with available resting-state data, the current study relied on an initial total of 316 sessions of imaging data acquired from 151 participants (79 females; age range 12–31) who participated in the first wave of data collection. Of the initial sample, 106 participants returned for a second visit (54 females; age range 13.5–33), and 59 participants returned for a third visit (26 females; age range 15–33). Participants were recruited from the community and were screened for the absence of psychiatric or neurological conditions, including loss of consciousness, for the absence of self-or first-degree relatives with reported major psychiatric illness, and for MRI contraindications (e.g., claustrophobia, metal in the body, pregnancy). Participants, or the parents of minors, gave informed consent, with those less than 18 years of age providing assent. All experimental procedures were approved by the University of Pittsburgh Institutional Review Board and complied with the Code of Ethics of the World Medical Association (Declaration of Helsinki, 1964).

### Imaging acquisition

MRI data were collected on a 3T Siemens Biograph mMR scanner, and structural, relaxometry, and resting-state fMRI sequences were acquired using the same protocol parameters across visits to ensure consistency in longitudinal measurements. Participants’ heads were stabilized with foam padding inside the head coil, and earbuds were provided to reduce scanner noise and improve comfort.

#### Structural acquisition and preprocessing

Structural images were acquired using a T1 weighted magnetization-prepared rapid gradient-echo (MPRAGE) sequence (TR, 2300 ms; echo time (TE), 2.98 ms; flip angle, 9°; inversion time (T1), 900 ms; voxel size, 1.0 × 1.0 × 1.0 mm). Structural MRI data were preprocessed with a pipeline that included skull-striping and warping to the MNI standard brain using both linear (FLIRT) and non-linear (FNIRT) transformations^71,72^.

#### Relaxometry data acquisition and preprocessing

Relaxometry data were acquired using multi-echo GRE (mGRE) and multi-echo turbo spin echo (mTSE) sequences to estimate R2* and R2 relaxation, respectively. The mGRE acquisition used: TE, 3, 8, 18, and 23 ms; TR, 724 ms; flip angle, 25°; FoV, 240 × 240 mm2. The mTSE used: effective TE, 12, 86, and 160 ms; TR, 6580 ms; 12 ms spacing between spin refocusing pulses; FoV, 240 × 240 mm2; 27 3 mm transverse slices; 1 mm slice gap. Quadratic penalized least squares (QPLS) was used to estimate R2* and R2 from the mGRE and mTSE magnitude images. R2’ was derived from the acquired R2 and R2* data for each participant by subtracting R2 from R2* estimates (see^38^ for a detailed description of estimate calculation methods). Prior to subtraction, R2 and R2* images were registered to MNI space using AFNI^73^. R2 registration involved concatenating the affine registration between the first echo of the mTSE and the anatomical image, and the non-linear registration of the anatomical image to MNI space. For the R2* registration, we added a rigid-body registration between the first echoes of the mGRE and mTSE images to the concatenation. Restricting the relaxometry data to sessions for which there was also resting-state fMRI data available resulted in 303 sessions of available R2* and R2’ data. Quality control procedures for all R2, R2*, and R2’ data have been described previously in^38^, and resulted in the exclusion of 67 sessions of R2* and R2’ data, for a total of 236 sessions of R2* and R2’ data initially included.

#### Resting-state fMRI data acquisition and preprocessing

Two 8-minute runs of fixation cross resting-state fMRI data were collected. Functional images were acquired using T2*-weighted signal from a single-echo, multiband EPI sequence (multiband factor = 3; TR, 1500 ms; TE, 30 ms; flip angle, 50°; voxel size, 2.3 × 2.3 mm in-plane resolution) with contiguous 2.3mm – thick slices aligned to maximally cover the cortex and basal ganglia. 316 sessions (with 628 runs) of resting-state data were initially included.

##### Preprocessing

Resting-state data were minimally preprocessed by performing simultaneous 4D slice-timing and motion correction^74^, skull stripping, intensity thresholding, wavelet despiking^75^, co-registration to the structural T1 and nonlinear warping to MNI space. Critically, minimal preprocessing of the resting-state fMRI data did not include operations that alter the absolute signal magnitude or remove global intensity trends (e.g., no temporal filtering, temporal normalization, nuisance regression, or temporal demeaning), as these standard fMRI steps are designed to isolate temporal fluctuations (BOLD) but would remove the cross-regional and inter-individual variance in the T2*w signal intensity that indexes tissue iron content.

### Derivation of candidate of T2*w brain iron measures

All T2*w measures were derived from the T2*w signal intensity of a single-echo EPI acquisition used in conventional fMRI studies (e.g., resting-state or task-based fMRI), requiring minimal additional processing (see **Figure 1** for a graphic depiction of the processing steps). Following the minimal preprocessing steps described above and the exclusion of high-motion volumes (framewise displacement ≥ 0.3 mm), the signal in each voxel (*v*) of a given volume was normalized to a reference value (*ref*), which was derived from a tissue reference mask (e.g., a whole-brain mask). These normalized voxel values were then averaged across all volumes to maximize the signal-to-noise ratio. To evaluate the robustness of the derived T2*w measures to differences in processing choices, we systematically evaluated multiple analytic permutations of each step in the processing pipeline, including: (1) reference tissue selection; (2) reference value aggregation; (3) the volume-wise normalization equation used to compute the T2*w measure; and (4) the voxelwise aggregation across volumes. These analytic options are further detailed below.

**Figure 1.**
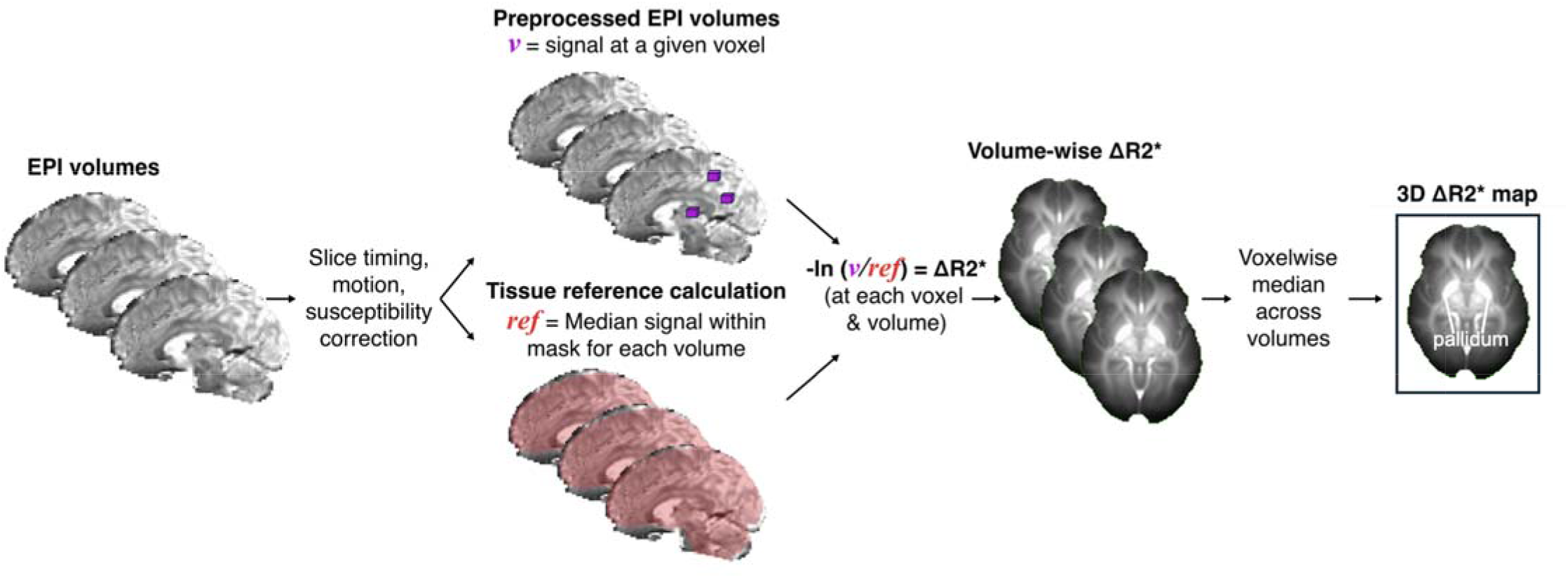
Processing steps for deriving ΔR2*. Delta R2* (ΔR2*) was derived from the T2*-weighted signal of a single-echo EPI acquisition used in conventional fMRI studies. The processing steps include standard preprocessing corrections for slice-timing (if applicable), motion, and susceptibility distortion; a signal referencing step in which the signal in each voxel of a given volume is normalized to a chosen reference value (e.g., a whole-brain average); and finally, to the enhance signal-to-noise ratio, the voxelwise ΔR2* signal is averaged over time (volumes) producing a 3D ΔR2* map. After comparing 43 possible combinations of pipeline configurations, we selected the following preferred set of processing steps depicted here: a tSNR-informed whole brain mask served as the tissue reference, the signal in each voxel was normalized according to Δ*R2*= −ln(v/ref)*, where *v* is the voxel intensity and *ref* is the reference value, and the ΔR2* signal intensity was averaged over volumes using the median. Note that the reference value is calculated separately for each volume to account for global changes in signal over time. Using this approach, the resulting ΔR2* signal is proportional to the difference in R2* between the signal in a given voxel and the reference tissue^68^, with higher signal intensity (greater brightness) in iron-rich areas (e.g., the pallidum).

#### 1. Defining reference tissue regions

Based on previous research on MRI-based iron quantification^66,76,77^, the following regions were initially chosen to serve as potential reference tissue regions: the corpus callosum, the ventricles, and the global signal of the whole brain. These reference tissue regions were defined as follows: the corpus callosum was defined using the body of corpus callosum region from the JHU ICBM-DTI-81 atlas^78^; the ventricles reference was created from a mask that included the first, second, and third ventricles as defined using the CerebrA atlas^79^; and a whole-brain reference region was defined using a coverage map created from all resting-state functional inputs for each participant. To reduce the influence of areas of poor signal quality due to signal dropout or other artifacts on the whole-brain reference value, we also created a tSNR-restricted whole-brain mask. First, a tSNR map was computed for each participant and thresholded at tSNR > 3 to identify voxels with adequate temporal stability. We then created a group-level consensus mask by selecting voxels that survived the tSNR threshold in >95% of sessions. This consensus mask was then applied to each volume to ensure that the reference value was derived from a consistent set of voxels across the cohort while systematically excluding regions prone to signal dropout and magnetic susceptibility artifacts.

#### 2. Reference tissue aggregation value calculation

For each reference region, the reference value was calculated separately for each volume to account for global changes in signal over time. Across all non-zero voxels in each reference region mask, we computed the average signal in order to aggregate the reference tissue signal into a single value, which served as the volume-wise reference value. We compared two aggregation methods, the mean and median.

#### 3. Volume-wise tissue reference normalization calculation

The signal intensity of each voxel within a volume was next normalized to the volume-specific reference tissue value. We evaluated four candidate equations motivated by prior work^80–82^:

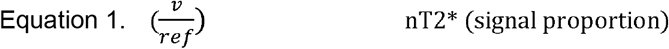

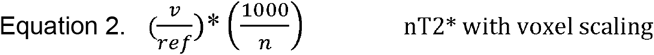

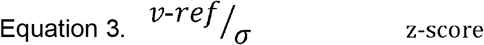

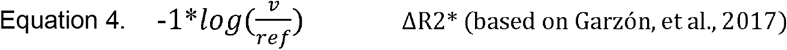

In these equations, *v* represents the voxel signal intensity, *ref* represents the aggregated reference value for the corresponding volume, and σ is the standard deviation of signal across the reference region. While Equations 1-3 have been used in past work to reflect relative signal intensity, they produce values that are inversely related to iron content, which presents a disadvantage for intuitive interpretation. Further, Equation 3 is most sensibly applied with a whole-brain reference, given that small or low-signal reference regions such as the corpus callosum or ventricles may exhibit near-zero or unstable standard deviations, leading to potentially unstable and difficult-to-interpret T2*w measure values when z-scoring is used. In contrast, Equation 4 applies a negative log-transformation to the signal ratio between the voxel and reference value that simplifies to the difference in the relaxation rate, R2* (R2*=1/T2*), between the voxel and reference value (ΔR2* = R2*_*v*_ – R2*_*ref*_), scaled by the echo time (TE). Thus, ΔR2* values scale positively with tissue iron concentration, aligning the directionality of the measure with established quantitative relaxometry (R2* and R2’) and facilitating more straightforward interpretation.

#### 4. Voxelwise averaging over volumes (temporal averaging)

Finally, to enhance signal-to-noise ratio, the voxelwise signal for the T2*w measures was averaged over volumes (time) using either the mean or median signal value, producing a static 3D map of the voxelwise T2*w measure for each participant.

#### Evaluation of processing pipelines

To evaluate the stability and robustness of the derived T2*w measures across different processing choices, we examined the T2*w measures from a total of 43 unique processing pipelines that differed in terms of 1) reference region used (corpus callosum, ventricles, whole brain, or a tSNR–informed whole-brain mask); 2) reference aggregation method (taking the mean vs. median of across all voxels within the reference region mask); 3) volume-wise tissue reference normalization calculations used to derive the T2*w measures (Eq. 1–4); and 4) the voxelwise method used to average the T2*w measure across volumes (taking the mean vs. median voxelwise signal across volumes).

### Defining basal ganglia regions of interest

Basal ganglia regions of interest (ROIs) were defined using the Harvard-Oxford Subcortical Atlas^83^. For each imaging measure of interest (ΔR2*, R2*, and R2′), the signal intensity was extracted from each lateralized basal ganglia ROI: the nucleus accumbens, caudate, pallidum, and putamen, and signal intensity was combined across the hemispheres to create bilateral ROIs. Additionally, signal from all bilateral ROIs was pooled to create an overall ‘all regions’ bilateral ROI (pooled across all voxels). Extracted values from this set of five bilateral ROIs served as inputs for all subsequent analyses.

### Correction for multiple comparisons

As all statistical testing was conducted across five ROIs, we applied the Benjamini–Hochberg false discovery rate (*FDR*) correction^84^ within each respective set of statistical tests, to account for multiple comparisons in significance testing across the five ROIs.

### T2*w measure quality control

Across all derived T2*w measure data, ten total runs of data were excluded for poor data quality including poor coverage for any region of interest or excessive signal variability (SD>6), reflecting excessive noise or artifacts within any of the basal ganglia regions of interest, resulting in 618 total runs of T2*w measure data across 316 sessions.

### Processing pipeline evaluation

#### Similarity across candidate pipelines

To evaluate how different processing pipelines affected the T2*w brain iron measures, we examined variation across measures derived from the different combinations of analytic options that comprised the 43 processing pipelines. First, absolute value Pearson’s correlations were computed between T2*w measures derived from each pairing of processing pipelines, across all participants, runs, and visits and all basal ganglia ROIs, producing 3,090 unique data points for each pipeline (618 total runs x 5 ROIs). Next, for each pipeline, we computed a ‘similarity score’ by averaging its correlation coefficients with all 42 other pipelines. Each pipeline was ranked by its similarity score to identify the most consistent and robust pipeline configuration.

#### Evaluating the influence of tissue reference

We looked at the effect of the tissue reference in two ways. First, we observed that reference region appeared to have the greatest impact on the T2*w measures across the processing pipelines (see **Supplementary Figure 1**). To quantify this, we grouped the pipelines by reference region and computed the mean similarity score for each group. We also compared the ΔR2* values derived using different reference regions to the relaxometry-based R2* and R2’ measures collected in the same study. To do so, we used linear mixed effects models (with a subject-level random intercept to account for repeated measures across runs and sessions) to compare the variance explained in both R2* and R2’ from the fixed effect of ΔR2* from analogous processing pipelines that differed only by reference region.

#### Selection of preferred pipeline

The final “preferred” processing pipeline was selected (the ΔR2* pipeline; see Results for full details of the selected pipeline) based on empirical results of the pipeline comparisons alongside practical considerations regarding signal stability and interpretability. This preferred pipeline was used for all subsequent analyses.

#### Model-based outlier detection

Prior to assessing validity and reliability, we performed model-based outlier detection on a per-ROI basis for ΔR2*, R2*, and R2’. Model-based outliers were identified as extreme values after accounting for well-known age-related differences in signal intensity. Using generalized additive mixed models (GAMMs), we estimated expected age-related variation in each measure and each ROI. GAMMs were fit using the *mgcv* library in R^85,86^. Specifically, within each basal ganglia ROI, we fit separate GAMMs for each imaging measure that modeled the effect of age as a smooth function (k = 6) using a thin-plate regression spline, and including fixed effects of visit number (as well as fixed effects of run in the case of ΔR2*) and subject-level random intercepts to account for repeated measures. Outliers, identified as residuals greater than three standard deviations beyond the predicted age-mean, were excluded from further analyses. This resulted in the exclusion of 18 out of 3090 total ΔR2* datapoints within the preferred ΔR2* processing pipeline. Out of the 1180 total relaxometry datapoints (236 sessions of relaxometry data x 5 ROIs), the ROI-level outlier removal resulted in the exclusion of 11 total datapoints for the R2* data, and 13 total datapoints for the R2’ data. The final sample sizes for each analysis for each ROI are provided in **Supplementary Table 1**.

#### Validation

We evaluated the validity of ΔR2* by examining its correspondence with R2* and R2’, well-established relaxometry-based indices of brain iron. R2* is an estimate of the rate of decay of T2* signal magnitude due to local magnetic-susceptibility inhomogeneities^68^ and served as the primary criterion measure. R2’ reflects the reversible transverse relaxation rate (1/T2’)^62,87^ and served as the secondary criterion measure. Following ROI-level outlier removal, validity analyses relied on approximately ~230 sessions of data per ROI (see **Supplementary Table 1** for precise samples for all ROIs and all analyses). In separate models, we regressed each relaxometry-based measure (R2* and R2’) on ΔR2* using a linear mixed-effects model that included a random intercept for participant to account for repeated measures within subject. We computed fixed effect *R*^2^ values from each model using the r2mlm package^88^ in R, to quantify, for each relaxometry-based measure (R2* or R2′), the proportion of variance explained by ΔR2*, with higher *R*^2^ values indicating stronger correspondence between ΔR2* and the established iron-sensitive measure.

#### Reliability and longitudinal stability

To evaluate the immediate test-retest reliability and longitudinal stability of ΔR2*, we conducted a series of analyses across repeated measures of ΔR2*, which was derived from two sessions of resting-state data collected at each of three longitudinal visits. First, to evaluate test-retest reliability for fMRI runs acquired in the same scanning session (~30-40 minutes apart; ~300 total pairs per ROI), we regressed ΔR2* values derived from the first resting-state run on the second resting-state run across all visits, separately for each ROI, using a linear mixed-effects model with a random intercept per subject to account for repeated measurements across study visits. To ascertain whether this relationship varied across longitudinal visits, we also calculated test-retest reliability separately for each longitudinal visit using ordinary least squares models. Test-retest reliability was estimated as the model *R*^2^, quantifying the proportion of shared variance between immediately repeated measurements of ΔR2*.

Next, we assessed the longitudinal stability of the ΔR2* measure across the three longitudinal visits (spaced ~1.5 years apart; >520 total visits). We calculated the intraclass correlation coefficients (ICCs) using linear mixed-effects models with a random intercept per subject and a fixed effect of visit number. The ICC reflects the proportion of total variance attributable to between-subject (trait-like) differences after accounting for the longitudinal effect of visit. ICC was calculated separately for each resting state fMRI run and additionally pooled over all runs and visits.

### Assessing the scalability of ΔR2* in the Adolescent Brain and Cognitive Development (ABCD) study

To demonstrate the scalability of the ΔR2* measure to large-scale imaging datasets, we derived ΔR2* using resting-state imaging data acquired at the baseline visit of the Adolescent Brain Cognitive Development (ABCD) Study.

### Sample 2 participants

The ABCD Study^61^ is the largest longitudinal investigation of adolescent brain development in the United States, which recruited over 11,000 youth across 21 sites. Written consent was obtained from all participants. The current analyses focused on the baseline sample (ages 9–11), relying on a subset of resting-state data that passed quality control^89^ and that had a sufficient number of low-motion volumes (see below for motion censoring details), the final sample included 8,366 participants (aged 9–11 years, 4,117 females).

#### Resting-state fMRI data acquisition and preprocessing

Youth underwent brain imaging across 21 sites and three different 3T scanner platforms: Siemens Prisma, General Electric 750 and Philips. Data collected from testing site 22 was excluded as there were only 27 datapoints available from this site. Comprehensive details on the ABCD Study design, scanning protocols, acquisition parameters, and preliminary quality control procedures have been previously described in detail^61,89^. Participants completed up to four 5-minute runs of resting-state T2*-weighted scans acquired in the axial plane using a single-echo EPI sequence. Critically, we used minimally processed data from the ABCC, processed using the ABCD-BIDS pipeline^90^, which is a modified version of the Human Connectome Project pipeline^91,92^, and included correction of gradient-nonlinearity-induced distortion, realignment of the timeseries to correct for subject motion (using a 6 DOF FLIRT registration of each frame to the single-band reference image), distortion correction, and transformation of functional images to standard MNI space. Each participant’s available resting-state data (up to four runs) were concatenated, along with their run-level motion parameters, and a new censor file was created to retain 300 randomly selected volumes of low-motion (framewise displacement threshold <.3) data for each participant. Only participants with 300 low-motion resting-state volumes retained were included in analyses. By standardizing the number of volumes used, we minimized potential biases introduced by varying amounts of data aggregated across participants.

### ΔR2* derivation

We derived ΔR2* from the ABCD baseline resting-sate data using the preferred pipeline selected in the pipeline evaluation phase. For each volume, we normalized the signal from each voxel to a reference value, defined as the median signal across all voxels of a tSNR-informed whole-brain mask, using Equation 4. The voxelwise signal was averaged over volumes using the median to yield a 3D ΔR2* map for each participant.

### Defining regions of interest

The same ROI creation procedure used in the prior dataset was applied here, yielding the same set of basal ganglia ROIs. Extracted ΔR2* values from the accumbens, caudate, pallidum, putamen, and an ‘all regions’ composite served as inputs for all subsequent analyses.

### Harmonization of ΔR2* data across ABCD sites and scanners

To account for potential site-specific differences in fMRI data acquisition prior to data aggregation, we performed image harmonization across ABCD acquisition sites following established protocols for developmental datasets^93^. Following computation and extraction of ΔR2* from the basal ganglia ROIs, we used an extension of neuroCombat^94^, called CovBat-GAM^94–97^, to statistically remove non-biological variation (and covariance) across data acquisition sites while preserving relevant biological features of interest (e.g., age, sex). While neuroCombat identifies and corrects for batch (e.g., site) effects, neuroCovBat additionally adjusts for covariates when they may be of interest, such as in the present analyses. We specifically used *the covfam* function from the *ComBatFamily* library^97^ in R and implemented a generalized additive model (GAM) approach to characterize potential non-linear effects, given past findings indicating non-linear developmental trajectories of brain tissue iron (e.g.,^30,33^). Harmonization covariates in the present analyses included ‘age’ as a smooth term (k = 3) and ‘sex’ assigned at birth as a parametric term. Our “batch” variable was ‘site’, accounting for the 21 ABCD acquisition sites.

### Examination of age effects in ΔR2*

For each ROI, we examined effects of age on ΔR2* using generalized additive models (GAMs) with penalized spline smoothing functions implemented using the *mgcv* library in R^86^. GAMs modeled the global effect of age as a penalized smooth function (k = 6) to capture age-related variation common across sites. Additionally, age-by-site interactions were modeled using by-factor smooths of age (k = 4), which estimate a separate smooth curve for each acquisition site using a shrinkage spline basis that allows uninformative site-specific curves to be penalized toward zero. Additionally, models included sex as a linear parametric effect. Site-specific smooths were plotted to depict variability in age-related trajectories across sites; however, inferential testing relied solely on the global smooth of age. Global effects of age were subjected to FDR-correction to account for multiple comparisons in significance testing across the five ROIs.

### Examination of sex effects in ΔR2*

For each ROI, sex differences in ΔR2* were evaluated using the parametric fixed-effect coefficient for sex from the previously fit GAMs (controlling for a smooth term of age). Statistical significance of the sex effect was assessed using the model-derived t-statistic, with FDR-correction applied to account for multiple comparisons across the five ROIs. For visualization and interpretability, estimated marginal means of ΔR2* for males and females were computed using the *emmeans* package^98^, evaluated at the sample mean age. These marginal means were calculated both by averaging over sites and separately within each site, allowing assessment of the consistency of sex-related differences in ΔR2* across data collection sites.

## RESULTS

### T2*w measures are robust to analytic choices and primarily influenced by reference tissue selection

Across all processing pipelines, the derived T2*w measures were highly convergent. Approximately 77% of the pipelines had a mean cross-pipeline correlation 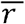 ≥ .85, demonstrating that the family of T2*w measures are highly robust to different analytic choices. Neither the reference aggregation or temporal averaging strategies (mean vs. median) substantially influenced the similarity scores. Instead, the choice of reference tissue had the largest impact on similarity of the derived T2*w measures (see **Supplementary Figure 1**). Pipeline configurations using the ventricle reference differed the most from the other reference regions (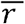 = .79). Pipelines using a corpus callosum (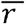 = .87) and a whole-brain reference region derived from the fMRI coverage map (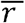 = .88) produced highly similar T2*w measures. The pipeline that used a tSNR-informed whole-brain reference region produced a T2*w measure that was most consistent with all other pipelines (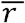 = .89).

### Selection of the ΔR2* reference normalization equation

While the reference normalization equations (Eq. 1–4) had minimal impact on similarity of the derived measures, Equation 4 (the ΔR2* calculation) was selected for its substantial interpretability advantages. Notably, the ΔR2* calculation is the only formulation that produces a T2*w measure that scales positively with tissue iron concentration and is directionally aligned with established R2* and R2’ measures.

### ΔR2* referenced to the tSNR-informed whole-brain mask shows the strongest convergence with quantitative relaxometry

Following the observation that reference region had a substantial impact on the T2*w measures, we explored which reference region produced a ΔR2* measure with the closest mapping to the gold-standard quantitative relaxometry measures collected from the same participants and scanning sessions. In a set of processing pipelines that held all analytic options constant except for the reference tissue, we compared the variance explained in the R2* and R2’ relaxometry-based measures by the respective ΔR2* measures. Across all basal ganglia regions, ΔR2* derived from the tSNR-informed whole-brain reference was most strongly associated with both R2* and R2’ (**Supplementary Figure 2**), indicating that ΔR2* referenced to the tSNR-informed whole-brain mask was most convergent with gold-standard iron metrics.

### The preferred ΔR2* processing pipeline

Based on the robustness and validity evaluations, alongside considerations of interpretability of the resulting measure, we ultimately selected a ΔR2* processing pipeline (as depicted in **Figure 1**) with the following features: 1) a tSNR-informed whole-brain reference region; 2) a reference tissue value derived from computing the median signal across all voxels in the reference region; 3) volume-wise tissue reference normalization using Equation 4 (the ΔR2* calculation); and 4) computing the median voxelwise signal across volumes (time) to enhance signal-to-noise in the final 3D ΔR2* map. Using this approach, the resulting ΔR2* value represents the magnitude of the signal intensity in a given voxel relative to the whole brain average such that higher values align with greater relative iron content. This ΔR2* processing pipeline was used for all subsequent analyses.

### ΔR2* aligns with quantitative relaxometry-based validation measures

Across the basal ganglia, the ΔR2* measure showed robust validity when evaluated against quantitative relaxometry-based metrics, which themselves have been validated against postmortem brain tissue iron content^99^. First, the derived ΔR2* measure exhibited spatial patterns that closely resembled those of R2* (**Figure 2a**) and R2′ (**Supplementary Figure 3a**), with the highest signal intensity observed in the iron-rich basal ganglia. Consistent with these spatial similarities, ΔR2* captured established regional differences in iron content across the basal ganglia for R2* (**Figure 2b**) and R2′ (**Supplementary Figure 3b**). Across all regions of the basal ganglia, ΔR2* was consistently associated with R2* (*R*^2^ accumbens = .39; *R*^2^ caudate = .05; *R*^2^ pallidum = .42; *R*^2^ putamen = .33; *R*^2^ all regions = .21; all *FDR* ≤ .001) and with R2′ (*R*^2^ accumbens = .40; *R*^2^ caudate = .01; *R*^2^ pallidum = .41; *R*^2^ putamen = .35; *R*^2^ all regions = .24; all *FDR* < .001, except caudate *FDR* = .234).

**Figure 2.**
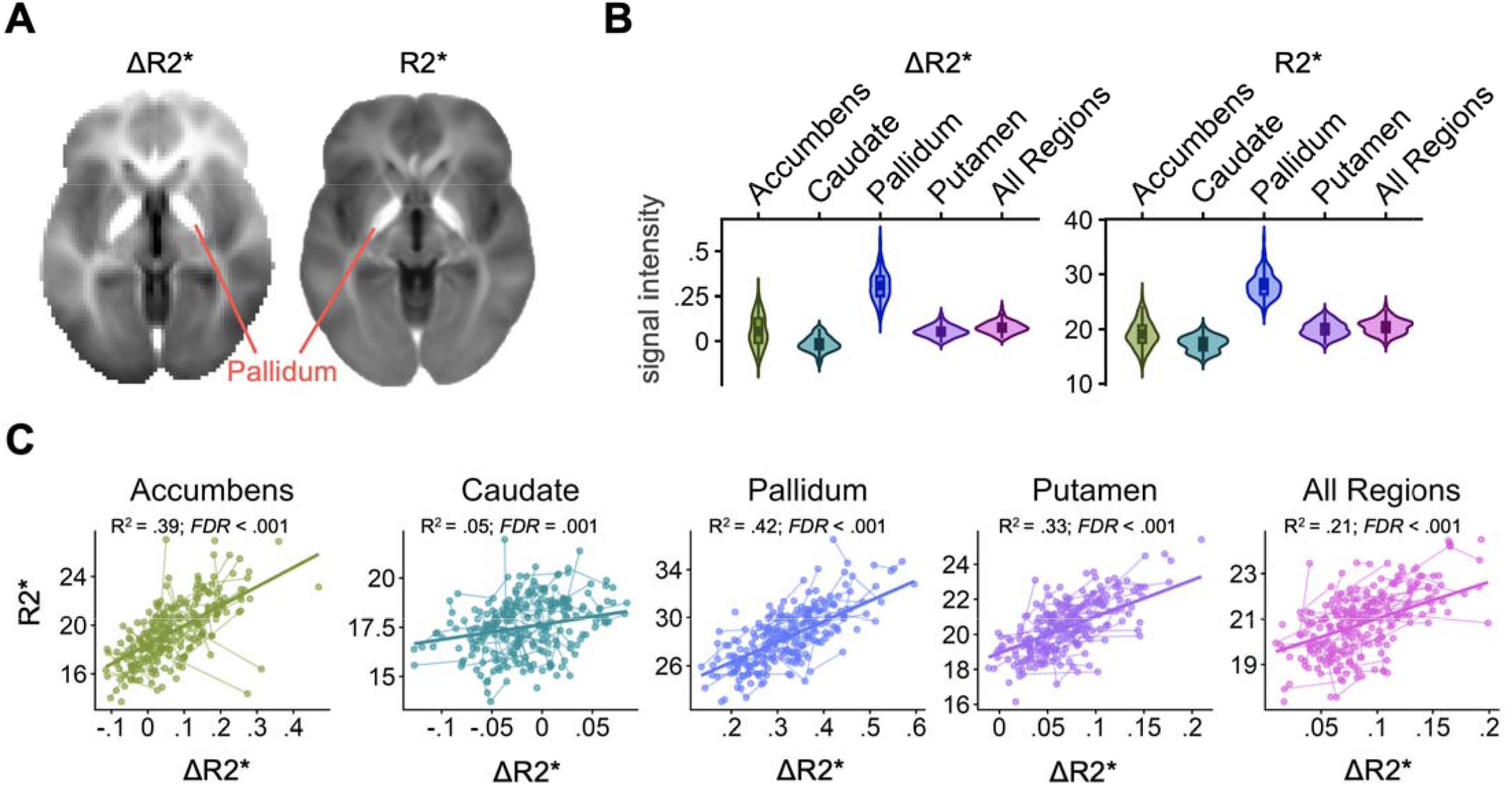
ΔR2* aligns with quantitative R2* relaxometry-based iron mapping across the basal ganglia. To establish the validity of the ΔR2*, we compared ΔR2* maps derived from resting-state fMRI data to R2* relaxometry-based iron maps in a sample that contained both measures. A) The ΔR2* and R2* measures showed similar spatial patterns with higher signal intensity (greater brightness) in iron-rich areas (e.g., the pallidum). B) ΔR2* and R2* measures show similar signal distributions and rank ordering across regions of the iron-rich basal ganglia, with the highe t signal in the pallidum. C) ΔR2* is consistently associated with R2* across regions of the basal ganglia (*FDR* ≤ .001), after outlier removal on a per region level: accumbens n = 229; caudate n = 232; pallidum n = 231; putamen n = 229; all regions n = 229.

### ΔR2* is highly reliable and longitudinally stable

Across all regions of the basal ganglia, ΔR2* demonstrated excellent reliability in terms of immediate test-retest reliability between fMRI runs acquired in the same scan session, longitudinal stability over visits (~1.5 years apart), and overall stability across runs and visits. Within-session comparisons of ΔR2* derived from resting-state fMRI runs collected in the same scan session showed excellent correspondence (**Figure 3a**), with immediate test–retest reliability *R*^2^ values across regions ranging from 0.79 to 0.95 within each visit and across all visits (**Figure 3b**). Longitudinal stability across the three study visits was also excellent with ICCs ranging from .69 to .84 across ROIs. The total ICCs across all runs and visits ranged from .75 to .85 across ROIs (**Figure 3c**), indicating that a substantial proportion of variance in ΔR2* reflected stable between-person differences over time.

**Figure 3.**
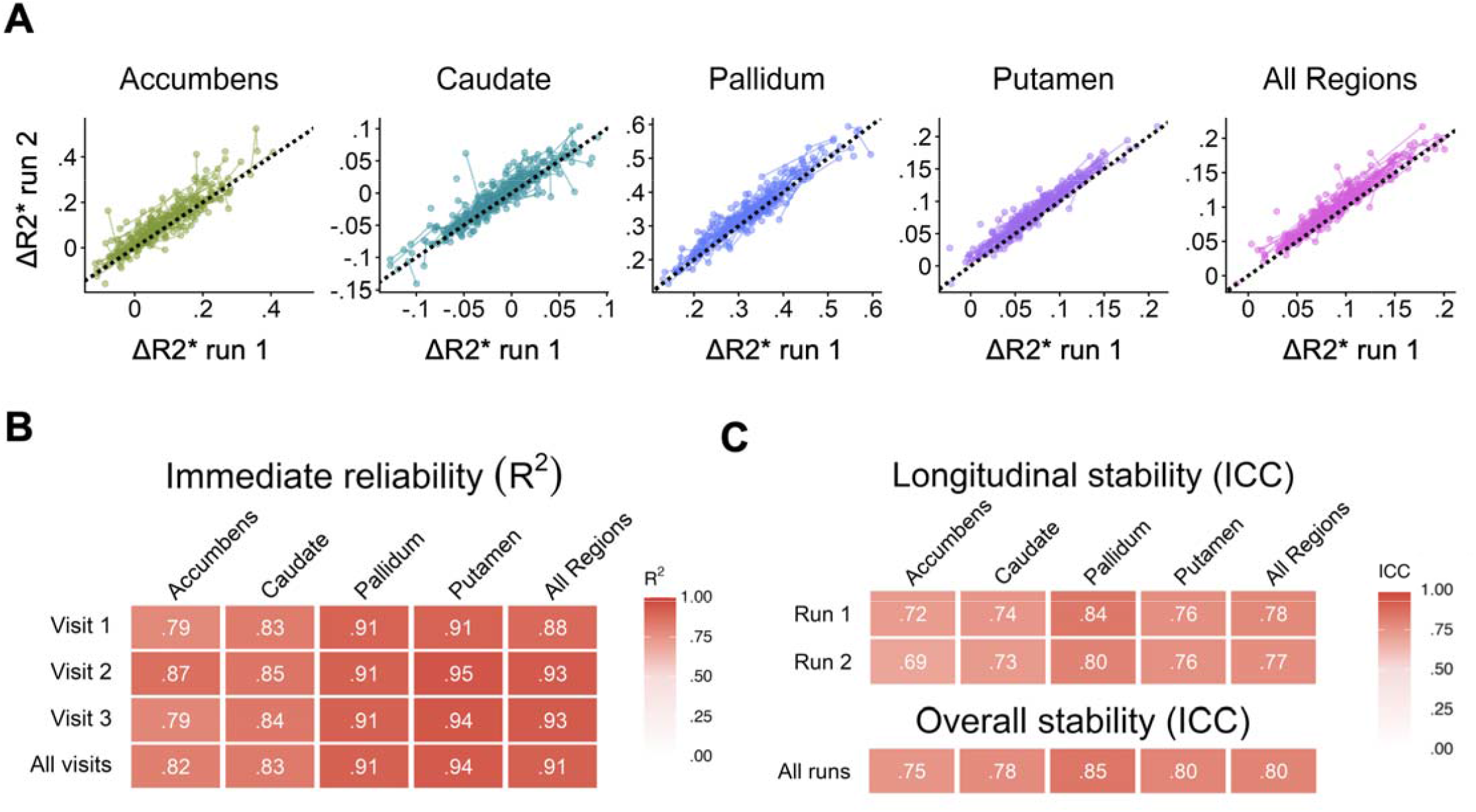
ΔR2* is highly reliable and longitudinally stable. To assess the reliability and stability of ΔR2* across repeated measures, we compared ΔR2* values from two fMRI runs acquired in the same session and across three longitudinal visits (~1.5 years apart). A) Across regions of the basal ganglia, ΔR2* showed excellent within-session correspondence between fMRI runs. B) Immediate test-retest reliability was high across all regions and visits (*R*^2^ = .79 - .95; *FDR* < .001). C) Longitudinal numeric stability was also high, with intraclass correlations (*ICC*) of .69 - .84 across visits within runs and .75 - .85 across the entire dataset.

### Large-scale application of ΔR2* captures known regional variation in iron content across the basal ganglia

We derived ΔR2* from resting state fMRI data acquired in 8,366 youth from the ABCD Study and harmonized values across the 21 ABCD collection sites using neuroCovBat-GAM. The resulting ΔR2* signal differences captured patterns that reflected both known differences in iron content in basal ganglia regions^99^ as well as patterns seen in the validation sample, including greater signal intensity in the pallidum and relatively high variability of signal in the accumbens (**Figure 4a and 4b**). While pre-harmonization ΔR2* values varied systematically across the 21 ABCD acquisition sites, harmonization effectively reduced site-related signal differences while preserving biologically meaningful variation (**Figure 4c**).

**Figure 4.**
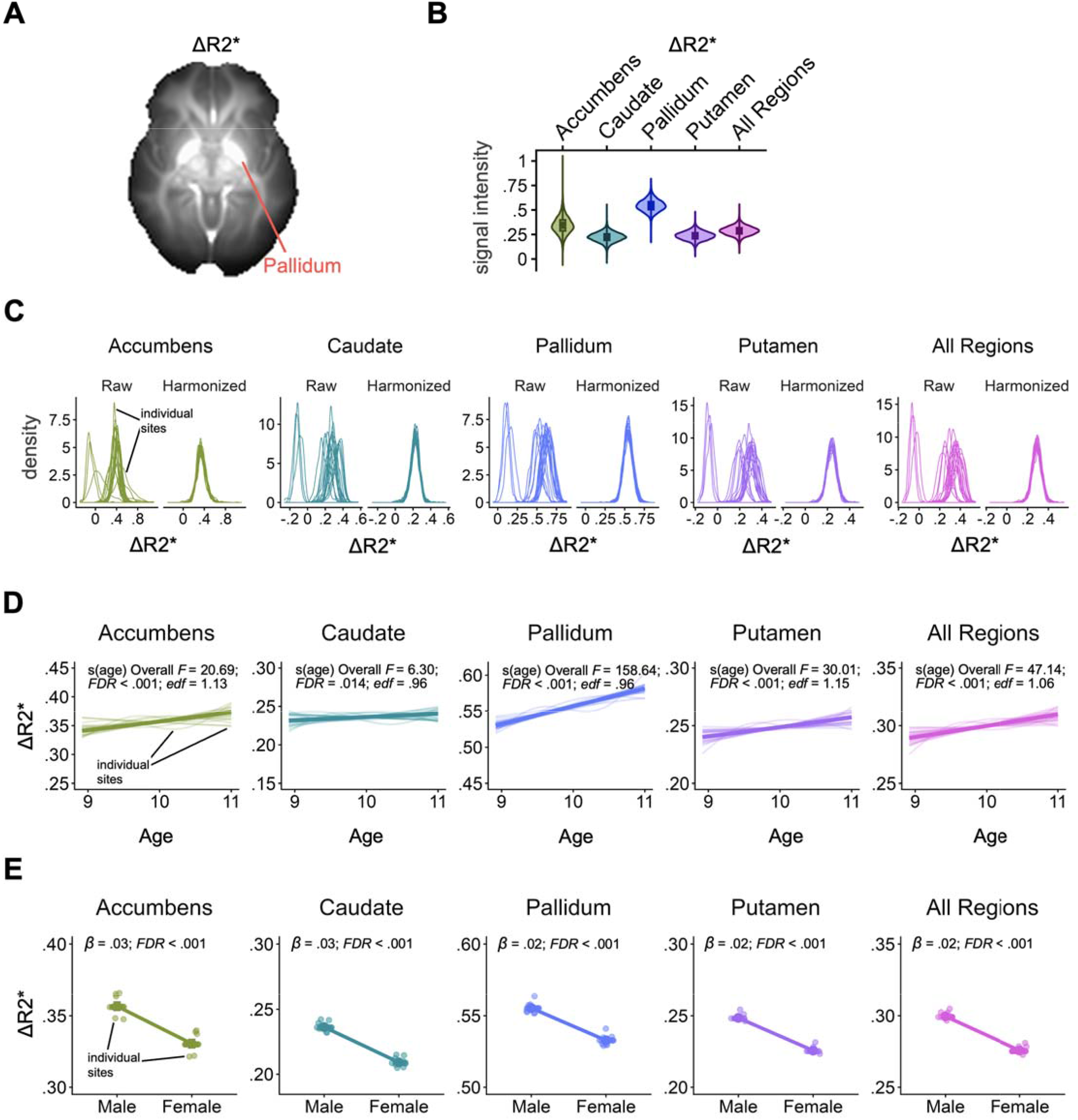
Large-scale application of ΔR2* in the ABCD Study: harmonization and age and sex differences. To demonstrate the scalability of the ΔR2* measure to existing large-scale datasets to allow retrospective estimation of brain iron differences, ΔR2* was derived from resting-state fMRI data in the baseline sample of the large scale ABCD Study (n = 8,366). A) The ABCD baseline-derived ΔR2* measure yielded a mean signal map with higher signal intensity (greater brightness) in iron-rich areas (e.g., the pallidum). B) ΔR2* showed expected variation in regional average signal across regions of the iron-rich basal ganglia, with the highest signal in the pallidum. C) Estimated ΔR2* values varied across the 21 ABCD sites (left) prior to harmonization. Site differences were successfully minimized (right) using harmonization procedures via neuroCovBat-GAM (Chen *et al*., 2021). D) Across all basal ganglia regions, ΔR2* increased with age from ages 9-11 years (all *FDR* ≤ .01), aligning with well-known age-related increases in brain iron content during this period^30–32,34,38,43^. E) ΔR2* was significantly higher in males in comparison to females (β = .02 - .03; *FDR* < .001), indicating sex-based variation in iron content.

### Basal ganglia ΔR2* increases with age across late childhood and is greater in males

Following data harmonization, ΔR2* replicated well-known age-related increases in brain iron content^30–32,34,38,43^ across all basal ganglia regions in ABCD baseline youth aged 9 to 11 years (accumbens: *edf* = 1.13, F = 20.69, *FDR* < .001; caudate: *edf* = .96, F = 6.30, *FDR* = .014; pallidum: *edf* = .96, F = 158.64, *FDR* < .001; putamen: *edf* = 1.15, F = 30.01, *FDR* < .001; all regions: *edf* = 1.06, F = 47.14, *FDR* < .001). In addition, significant sex differences were observed across all regions, with males showing higher ΔR2* values than females (accumbens: β = 0.026, t = 13.42, *FDR* < .001; caudate: β = 0.027, t = 26.13, *FDR* < .001; pallidum: β = 0.023, t = 17.87, *FDR* < .001; putamen: β = 0.023, t = 24.06, *FDR* < .001; all regions: β = 0.024, t = 24.65, *FDR* < .001).

## DISCUSSION

We establish delta R2* (ΔR2*) as a reliable, scalable, and developmentally sensitive marker of basal ganglia tissue iron that can be derived from standard single-echo echo-planar imaging (EPI) acquired during conventional fMRI. We demonstrate that ΔR2* converges with R2* and R2’– quantitative relaxometry measures with post-mortem validation^25,58^ – and exhibits both high short-term reliability and robust longitudinal stability across timescales ranging from hours to years. Importantly, ΔR2* was effectively scaled to the Adolescent Brain and Cognitive Development (ABCD) cohort^61^, where it was sensitive to established developmental and sex-linked variation during adolescence, demonstrating its scalability and utility in large, population-scale datasets lacking dedicated iron imaging. Beyond validation, we provide practical harmonization strategies and processing recommendations alongside a Brain Imaging Data structure (BIDS) application for investigators to readily derive ΔR2* in new and existing datasets. Across 43 processing pipelines, ΔR2* showed strong robustness to analytic and design variations, positioning it as a low-burden, widely accessible tool for retrospective and prospective studies. Together, these findings substantially expand access to iron-sensitive non-invasive neurobiological markers for developmental and translational neuroscience.

### Design Considerations and Validation

ΔR2* estimates the relative difference in R2* between the target tissue (e.g., basal ganglia) and a reference region using within-scan normalization, enhancing sensitivity to iron-related signal variation^68^. Although the computational framework is straightforward, several analytic decisions could plausibly influence the stability and interpretability of ΔR2* estimates. Accordingly, we systematically evaluated four core design components: (1) reference tissue selection; (2) reference value aggregation (mean vs. median); (3) the normalization equation used to compute ΔR2* within each volume; and (4) voxelwise aggregation across volumes (mean vs. median). We demonstrate that ΔR2* is highly robust to most analytic decisions, with high convergence across pipelines, indicating a significant amount of analytic flexibility. Notably, reference tissue selection emerged as a dominant source of variation in ΔR2*, with pipelines using ventricular reference regions exhibiting substantially greater variability, whereas whole brain and corpus callosum reference strategies, as used in prior work^66,76,77^, yielded highly similar estimates. Here, we found that a tSNR-informed whole-brain reference region minimized variability across pipeline configurations, provided the most stable estimates, and maximized correspondence with quantitative relaxometry measures (R2* and R2’), establishing it as both the most analytically stable and biologically valid choice in this developmental sample. This may, in part, reflect the use of a developmental sample in which global brain properties – such as total brain volume^100^ – are still undergoing maturation, where a whole-brain reference may provide a robust normalization strategy by averaging across widespread developmental variation. However, the optimal reference region may depend on the population under study, and alternative reference strategies (e.g., white matter-restricted or corpus callosum-based references) should be empirically tested within each study’s specific population and acquisition context, rather than assuming a single optimal strategy.

In contrast, voxelwise and reference aggregation strategies (mean vs. median) exerted minimal influence on ΔR2* similarity. Nevertheless, we recommend median-based aggregation as it provides additional robustness to outlier volumes, an important consideration in developmental samples with higher motion. While our past work has used time-averaged and normalized T2* (nT2*) (Eq. 1 & 2), the signal proportion between the tissue and the reference, we now recommend a negative log-transformation normalization (as implemented by by Garzón et al.^68^), which we refer to as ΔR2* (Eq 4). This approach is preferred because it intuitively mathematically simplifies to the *difference* (rather than *proportion*) in R2* between the tissue and reference. Thus, the ΔR2* method produces values that scale positively with tissue iron concentration, aligning directionality with established R2* and R2’ measures and simplifying interpretation (relative to the inverse relationship between the T2* signal and iron content). Together, these stepwise analyses provide empirically grounded design recommendations that support ΔR2* as a robust and conceptually straightforward metric. Finally, we emphasize that fMRI data used for these calculations must be minimally preprocessed; specifically, researchers should avoid temporal filtering, mean-centering, or global signal normalization, as these standard BOLD-centric operations remove the absolute signal intensity variations that are essential for indexing underlying tissue iron content.

For iron metrics to support developmental and translational research, they should demonstrate longitudinal stability and permit reliable cross-site and cross-scanner comparison, particularly in longitudinal studies seeking to dissociate normative age-related change from trait-level differences and emerging psychiatric risk. Using the optimized pipeline, we conducted a comprehensive set of analyses testing validation, reliability, and scalability to evaluate whether ΔR2* functions as a stable marker of basal ganglia iron. Across basal ganglia ROIs, ΔR2* showed strong correspondence with R2* and R2’ and replicated well-established regional patterns of iron content^30–32^. ΔR2* also demonstrated excellent measurement reliability, with high immediate test-retest reliability across resting-state runs within the same scan session, as well as excellent longitudinal stability across visits spanning up to three years, indicating that a substantial proportion of variance reflects stable between-person differences rather than measurement noise. These properties are particularly important for developmental and longitudinal study designs aimed at disentangling trait-level individual differences from within-person change. Critically, we demonstrate the scalability of ΔR2* by applying this pipeline to the ABCD baseline cohort^61^ – a large, multi-site developmental dataset lacking dedicated iron imaging, similar to most human neuroimaging studies. After harmonizing to account for site-related variability, ΔR2* retained biologically meaningful signal, including age- and sex-effects, while substantially reducing non-biological effects of study site. Post-harmonization, ΔR2* distributions across ROIs replicated known patterns of iron levels in the basal ganglia. Further, across all basal ganglia ROIs, there were robust age-related increases in ΔR2*, even within the narrow baseline age range of 9-11 years, highlighting the sensitivity of this measure to subtle developmental change. Replication of sex differences in iron-related signal^35^ further support its biological validity. Collectively, these findings establish ΔR2* as a reliable, scalable, and biologically sensitive non-invasive approach for estimating basal ganglia iron in large-scale developmental neuroimaging studies, suggesting broad utility for developmental neuroscience.

### Limitations and Future Considerations

ΔR2* reflects relative differences in R2* between target and reference tissue iron rather than an absolute quantitative estimate, and as such, can be influenced by acquisition parameters such as echo time (TE), scanner-specific factors, including scanner upgrades and version, and time (longitudinal assessments) that can potentially limit cross-study comparability by introducing unwanted variance in iron sensitive signals. Without harmonization^96,101^, such variability can obscure developmental effects, inflate between-site differences, and limit clinical inference. Our harmonization analyses demonstrate that site effects on individual differences can be mitigated when acquisition protocols are well-matched, as in ABCD. However, careful inspection and harmonization remain essential, particularly in multi-site and longitudinal studies, or for mega-or meta-analyses. Future work should evaluate these impacts on ΔR2* and harmonization strategies for reducing non-biological sources of variation. Additionally, correlations between ΔR2* and established relaxometry-based iron markers were strongest in the NAcc and pallidum, regions with higher iron concentrations and stronger signal^28,102^. Correspondence was weaker in the caudate, likely reflecting its lower iron content reported in postmortem work^103^, as well as previously noted weaker correlations between *in vivo* imaging and postmortem iron measures in this region^102^. Notably, caudate ΔR2* showed similarly high reliability and longitudinal stability, values within the expected range for this region, and age-related effects consistent with prior literature^29,35,104^. Reduced correspondence with relaxometry measures may therefore reflect both biological factors (lower iron content, potential contributions from non-iron sources such as myelin) as well as methodological considerations including partial volume effects due to its proximity to ventricles. Although not directly tested here, prior work suggests that ΔR2* can be derived from both resting-state and task-based EPI^55^, and from single-or multi-echo acquisitions, increasing flexibility for retrospective analyses and reducing the burden of a dedicated acquisition sequence that can add time to prospective studies. Future work should characterize sensitivity across acquisition parameters and populations, including aging and clinical samples. Finally, as ΔR2* is not quantitative, it reflects an indirect measure of tissue iron concentration that has been linked to DA metabolism in humans^38^. As such, ΔR2* may be best viewed as a scalable discovery tool that can flag biologically meaningful variation warranting follow-up with more direct iron quantification methods (e.g., Quantitative Susceptibility Mapping (QSM), R2*, and R2’).

### Open Science and Community Impact

To facilitate broad adoption, we have developed a lightweight, publicly available processing tool that enables derivation of ΔR2* from standard fMRI datasets (https://zenodo.org/records/19475933). The workflow is flexible to design considerations (i.e., choice of a reference region and aggregation strategies), is compatible with BIDS-formatted data and integrates seamlessly with common preprocessing pipelines such as *fMRIPrep*^105^, enabling researchers to extract iron-sensitive metrics from both prospective and legacy datasets. By lowering methodological barriers, this resource positions the field to incorporate iron-sensitive neurobiology into developmental, clinical, and translational research at scale, facilitating mechanistic investigations of critical neurodevelopmental processes and potential pathways of clinical risk across the lifespan. By enabling iron-sensitive analyses in large, existing population-based legacy datasets, as well as prospective studies, ΔR2* supports prospective tests of whether accelerated, delayed, or exaggerated iron trajectories confer risk or resilience across decades. In this fashion, ΔR2* bridges developmental and aging research, facilitating integrative, transdiagnostic frameworks of basal ganglia iron as a brain-based marker that shapes normative development, psychiatric risk, and neurodegenerative processes across the lifespan.

## Acknowledgements

We thank the participants, the ABCD Study, and Laboratory of Neurocognitive Development (LNCD) research assistants.

## REFERENCES

1. Larsen, B. & Luna, B. Adolescence as a neurobiological critical period for the development of higher-order cognition. Neurosci. Biobehav. Rev. 94, 179–195 (2018).

2. Tervo-Clemmens, B. et al. A Canonical Trajectory of Executive Function Maturation During the Transition from Adolescence to Adulthood. Preprint at 10.31234/osf.io/73yfv (2022).

3. Spear, L. P. The adolescent brain and age-related behavioral manifestations. Neurosci. Biobehav. Rev. 24, 417–463 (2000).

4. Trifilieff, P. & Martinez, D. Blunted Dopamine Release as a Biomarker for Vulnerability for Substance Use Disorders. Biol. Psychiatry 76, 4–5 (2014).

5. Tervo-Clemmens, B., Quach, A., Calabro, F. J., Foran, W. & Luna, B. Meta-analysis and review of functional neuroimaging differences underlying adolescent vulnerability to substance use. NeuroImage 209, 116476 (2020).

6. Howes, O. D. & Shatalina, E. Integrating the Neurodevelopmental and Dopamine Hypotheses of Schizophrenia and the Role of Cortical Excitation-Inhibition Balance. Biol. Psychiatry 92, 501–513 (2022).

7. Larsen, B., Verstynen, T. D., Yeh, F.-C. & Luna, B. Developmental Changes in the Integration of Affective and Cognitive Corticostriatal Pathways are Associated with Reward-Driven Behavior. Cereb. Cortex 28, 2834–2845 (2018).

8. Wahlstrom, D., White, T. & Luciana, M. Neurobehavioral evidence for changes in dopamine system activity during adolescence. Neurosci. Biobehav. Rev. 34, 631–648 (2010).

9. Brenhouse, H. C. & Andersen, S. L. Developmental trajectories during adolescence in males and females: A cross-species understanding of underlying brain changes. Neurosci. Biobehav. Rev. 35, 1687–1703 (2011).

10. Andersen, S. L., Dumont, N. L. & Teicher, M. H. Developmental differences in dopamine synthesis inhibition by (±)-7-OH-DPAT. Naunyn. Schmiedebergs Arch. Pharmacol. 356, 173–181 (1997).

11. Luciana, M., Wahlstrom, D., Porter, J. N. & Collins, P. F. Dopaminergic modulation of incentive motivation in adolescence: age-related changes in signaling, individual differences, and implications for the development of self-regulation. Dev. Psychol. 48, 844–861 (2012).

12. Mitchell, M. R. et al. Adolescent Risk Taking, Cocaine Self-Administration, and Striatal Dopamine Signaling. Neuropsychopharmacology 39, 955–962 (2014).

13. Abi-Dargham, A. & Horga, G. The search for imaging biomarkers in psychiatric disorders. Nat. Med. 22, 1248–1255 (2016).

14. Brass, S. D., Chen, N., Mulkern, R. V. & Bakshi, R. Magnetic Resonance Imaging of Iron Deposition in Neurological Disorders. Top. Magn. Reson. Imaging 17, 31–40 (2006).

15. Connor, J. R., Menzies, S. L., Martin, S. M. S. & Mufson, E. J. Cellular distribution of transferrin, ferritin, and iron in normal and aged human brains. J. Neurosci. Res. 27, 595–611 (1990).

16. Morris, C. M., Candy, J. M., Oakley, A. E., Bloxham, C. A. & Edwardson, J. A. Histochemical Distribution of Non-Haem Iron in the Human Brain. Cells Tissues Organs 144, 235–257 (1992).

17. Thomas, L. O., Boyko, O. B., Anthony, D. C. & Burger, P. C. MR detection of brain iron. Am. J. Neuroradiol. 14, 1043–1048 (1993).

18. Ortega, R., Cloetens, P., Devès, G., Carmona, A. & Bohic, S. Iron Storage within Dopamine Neurovesicles Revealed by Chemical Nano-Imaging. PLoS ONE 2, (2007).

19. Zucca, F. A. et al. Interactions of iron, dopamine and neuromelanin pathways in brain aging and Parkinson’s disease. Prog. Neurobiol. 155, 96–119 (2017).

20. Ward, R. J., Zucca, F. A., Duyn, J. H., Crichton, R. R. & Zecca, L. The role of iron in brain ageing and neurodegenerative disorders. Lancet Neurol. 13, 1045–1060 (2014).

21. Connor, J. R. & Menzies, S. L. Cellular management of iron in the brain. J. Neurol. Sci. 134 Suppl, 33–44 (1995).

22. Todorich, B., Pasquini, J. M., Garcia, C. I., Paez, P. M. & Connor, J. R. Oligodendrocytes and myelination: The role of iron. Glia 57, 467–478 (2009).

23. Paul, B. T., Manz, D. H., Torti, F. M. & Torti, S. V. Mitochondria and Iron: current questions. Expert Rev. Hematol. 10, 65–79 (2017).

24. Möller, H. E. et al. Iron, Myelin, and the Brain: Neuroimaging Meets Neurobiology. Trends Neurosci. 42, 384–401 (2019).

25. Stüber, C. et al. Myelin and iron concentration in the human brain: A quantitative study of MRI contrast. NeuroImage 93, 95–106 (2014).

26. Stowell, R. & Wang, K. H. Dopaminergic signaling regulates microglial surveillance and adolescent plasticity in the mouse frontal cortex. Nat. Commun. 16, 7974 (2025).

27. Kim, Y. S., Choi, J. & Yoon, B.-E. Neuron-Glia Interactions in Neurodevelopmental Disorders. Cells 9, 2176 (2020).

28. Hallgren, B. & Sourander, P. The Effect of Age on the Non-Haemin Iron in the Human Brain. J. Neurochem. 3, 41–51 (1958).

29. Aquino, D. et al. Age-related Iron Deposition in the Basal Ganglia: Quantitative Analysis in Healthy Subjects. Radiology 252, 165–172 (2009).

30. Larsen, B. et al. Longitudinal Development of Brain Iron Is Linked to Cognition in Youth. J. Neurosci. 40, 1810–1818 (2020).

31. Larsen, B. & Luna, B. In vivo evidence of neurophysiological maturation of the human adolescent striatum. Dev. Cogn. Neurosci. 12, 74–85 (2015).

32. Peterson, E. T. et al. Distribution of brain iron accrual in adolescence: Evidence from cross-sectional and longitudinal analysis. Hum. Brain Mapp. 40, 1480–1495 (2019).

33. Parr, A. C. et al. Contributions of dopamine-related basal ganglia neurophysiology to the developmental effects of incentives on inhibitory control. Dev. Cogn. Neurosci. 54, 101100 (2022).

34. Hect, J. L., Daugherty, A. M., Hermez, K. M. & Thomason, M. E. Developmental variation in regional brain iron and its relation to cognitive functions in childhood. Dev. Cogn. Neurosci. 34, 18–26 (2018).

35. Parr, A. C. et al. Developmental variation in dopamine neurobiology, neurocognitive functioning, and impulsivity shape substance use trajectories in youth. 2025.10.24.684358 Preprint at 10.1101/2025.10.24.684358 (2025).

36. Daugherty, A. M. & Raz, N. Appraising the Role of Iron in Brain Aging and Cognition: Promises and Limitations of MRI Methods. Neuropsychol. Rev. 25, 272–287 (2015).

37. Ward, R. J., Zucca, F. A., Duyn, J. H., Crichton, R. R. & Zecca, L. The role of iron in brain ageing and neurodegenerative disorders. Lancet Neurol. 13, 1045–1060 (2014).

38. Larsen, B. et al. Maturation of the human striatal dopamine system revealed by PET and quantitative MRI. Nat. Commun. 11, 1–10 (2020).

39. Corrigan, N. M. et al. Myelin development in cerebral gray and white matter during adolescence and late childhood. NeuroImage 227, 117678 (2021).

40. Sydnor, V. J. et al. Heterochronous laminar maturation in the human prefrontal cortex. 2025.01.30.635751 Preprint at 10.1101/2025.01.30.635751 (2025).

41. Caballero, A., Orozco, A. & Tseng, K. Y. Developmental regulation of excitatory-inhibitory synaptic balance in the prefrontal cortex during adolescence. Semin. Cell Dev. Biol. 118, 60–63 (2021).

42. Larsen, B. et al. A Developmental Reduction of the Excitation:Inhibition Ratio in Association Cortex during Adolescence. http://biorxiv.org/lookup/doi/10.1101/2021.05.19.444703 (2021) doi:10.1101/2021.05.19.444703.

43. Parr, A. C. et al. Dopamine-related striatal neurophysiology is associated with specialization of frontostriatal reward circuitry through adolescence. Prog. Neurobiol. 201, 101997 (2021).

44. Christakou, A., Brammer, M. & Rubia, K. Maturation of limbic corticostriatal activation and connectivity associated with developmental changes in temporal discounting. NeuroImage 54, 1344–1354 (2011).

45. Sydnor, V. J. et al. Human thalamocortical structural connectivity develops in line with a hierarchical axis of cortical plasticity. Nat. Neurosci. 28, 1772–1786 (2025).

46. Hoops, D. & Flores, C. Making Dopamine Connections in Adolescence. Trends Neurosci. 40, 709–719 (2017).

47. Larsen, B. et al. Development of Iron Status Measures during Youth: Associations with Sex, Neighborhood Socioeconomic Status, Cognitive Performance, and Brain Structure. Am. J. Clin. Nutr. 118, 121–131 (2023).

48. Lozoff, B. Early Iron Deficiency Has Brain and Behavior Effects Consistent with Dopaminergic Dysfunction. J. Nutr. 141, 740S–746S (2011).

49. Lozoff, B. et al. Long-Lasting Neural and Behavioral Effects of Iron Deficiency in Infancy. Nutr. Rev. 64, S34–S43 (2006).

50. Barbosa, J. H. O. et al. Quantifying brain iron deposition in patients with Parkinson’s disease using quantitative susceptibility mapping, R2 and R2*. Magn. Reson. Imaging 33, 559–565 (2015).

51. Connor, J. R. et al. Profile of altered brain iron acquisition in restless legs syndrome. Brain 134, 959–968 (2011).

52. Adisetiyo, V. et al. Multimodal MR imaging of brain iron in attention deficit hyperactivity disorder: a noninvasive biomarker that responds to psychostimulant treatment? Radiology 272, 524–532 (2014).

53. Cascone, A. D. et al. Brain tissue iron neurophysiology and its relationship with the cognitive effects of dopaminergic modulation in children with and without ADHD. Dev. Cogn. Neurosci. 63, 101274 (2023).

54. Yao, S. et al. Quantitative Susceptibility Mapping Reveals an Association between Brain Iron Load and Depression Severity. Front. Hum. Neurosci. 0, (2017).

55. Price, R. B. et al. Biobehavioral correlates of an fMRI index of striatal tissue iron in depressed patients. Transl. Psychiatry 11, 1–8 (2021).

56. Sonnenschein, S. F. et al. Subcortical brain iron deposition in individuals with schizophrenia. J. Psychiatr. Res. 151, 272–278 (2022).

57. Ersche, K. D. et al. Disrupted iron regulation in the brain and periphery in cocaine addiction. Transl. Psychiatry 7, e1040 (2017).

58. Langkammer, C. et al. Quantitative susceptibility mapping (QSM) as a means to measure brain iron? A post mortem validation study. NeuroImage 62, 1593–1599 (2012).

59. Kilbourn, M. R. Radioligands for Imaging Vesicular Monoamine Transporters. in PET and SPECT of Neurobiological Systems (eds Dierckx, R. A. J. O., Otte, A., de Vries, E. F. J., van Waarde, A. & Luiten, P. G. M.) 765–790 (Springer, Berlin, Heidelberg, 2014). doi:10.1007/978-3-642-42014-6_27.

60. Wahlstrom, D., Collins, P., White, T. & Luciana, M. Developmental changes in dopamine neurotransmission in adolescence: behavioral implications and issues in assessment. Brain Cogn. 72, 146–159 (2010).

61. Casey, B. J. et al. The Adolescent Brain Cognitive Development (ABCD) study: Imaging acquisition across 21 sites. Dev. Cogn. Neurosci. 32, 43–54 (2018).

62. Haacke, E. M. et al. Imaging iron stores in the brain using magnetic resonance imaging. Magn. Reson. Imaging 23, 1–25 (2005).

63. Aoki, S. et al. Normal deposition of brain iron in childhood and adolescence: MR imaging at 1.5 T. Radiology 172, 381–385 (1989).

64. He, X. & Yablonskiy, D. A. Biophysical mechanisms of phase contrast in gradient echo MRI. Proc. Natl. Acad. Sci. 106, 13558–13563 (2009).

65. Chavhan, G. B., Babyn, P. S., Thomas, B., Shroff, M. M. & Haacke, E. M. Principles, Techniques, and Applications of T2*-based MR Imaging and Its Special Applications. Radiographics 29, 1433–1449 (2009).

66. Garzón, B., Sitnikov, R., Bäckman, L. & Kalpouzos, G. Can transverse relaxation rates in deep gray matter be approximated from functional and T2-weighted FLAIR scans for relative brain iron quantification? Magn. Reson. Imaging 40, 75–82 (2017).

67. Gandon, Y. et al. Non-invasive assessment of hepatic iron stores by MRI. The Lancet 363, 357–362 (2004).

68. Garzón, B., Sitnikov, R., Bäckman, L. & Kalpouzos, G. Can transverse relaxation rates in deep gray matter be approximated from functional and T2-weighted FLAIR scans for relative brain iron quantification? Magn. Reson. Imaging 40, 75–82 (2017).

69. Lehéricy, S., Bardinet, E., Poupon, C., Vidailhet, M. & François, C. 7 Tesla magnetic resonance imaging: a closer look at substantia nigra anatomy in Parkinson’s disease. Mov. Disord. Off. J. Mov. Disord. Soc. 29, 1574–1581 (2014).

70. Calabro, F. J. et al. Striatal dopamine supports reward expectation and learning: A simultaneous PET/fMRI study. NeuroImage 267, 119831 (2023).

71. Jenkinson, M., Beckmann, C. F., Behrens, T. E. J., Woolrich, M. W. & Smith, S. M. FSL. NeuroImage 62, 782–790 (2012).

72. Foran, W. et al. LabNeuroCogDevel/fmri_processing_scripts: init. Zenodo 10.5281/zenodo.8320245 (2023).

73. Cox, R. W. AFNI: software for analysis and visualization of functional magnetic resonance neuroimages. Comput. Biomed. Res. 29, 162–173 (1996).

74. Hallquist, M. N., Hwang, K. & Luna, B. The nuisance of nuisance regression: spectral misspecification in a common approach to resting-state fMRI preprocessing reintroduces noise and obscures functional connectivity. NeuroImage 82, 208–225 (2013).

75. Patel, A. X. et al. A wavelet method for modeling and despiking motion artifacts from resting-state fMRI time series. NeuroImage 95, 287–304 (2014).

76. QSM Consensus Organization Committee et al. Recommended implementation of quantitative susceptibility mapping for clinical research in the brain: A consensus of the ISMRM electro□magnetic tissue properties study group. Magn. Reson. Med. 91, 1834–1862 (2024).

77. Peterson, E. T. et al. Distribution of brain iron accrual in adolescence: Evidence from cross□sectional and longitudinal analysis. Hum. Brain Mapp. 40, 1480–1495 (2019).

78. Mori, S., Wakana, S., Nagae-Poetscher, L. M. & VanZijl, P. C. M. MRI Atlas of Human White Matter. vol. 1 (Elsevier B. V., Amsterdam, 2005).

79. Manera, A. L., Dadar, M., Fonov, V. & Collins, D. L. CerebrA, registration and manual label correction of Mindboggle-101 atlas for MNI-ICBM152 template. Sci. Data 7, 237 (2020).

80. Larsen, B. & Luna, B. In vivo evidence of neurophysiological maturation of the human adolescent striatum. Dev. Cogn. Neurosci. 12, 74–85 (2015).

81. Parr, A. C. et al. Contributions of dopamine-related basal ganglia neurophysiology to the developmental effects of incentives on inhibitory control. Dev. Cogn. Neurosci. 54, 101100 (2022).

82. Garzón, B., Sitnikov, R., Bäckman, L. & Kalpouzos, G. Can transverse relaxation rates in deep gray matter be approximated from functional and T2-weighted FLAIR scans for relative brain iron quantification? Magn. Reson. Imaging 40, 75–82 (2017).

83. Jenkinson, M., Beckmann, C. F., Behrens, T. E., Woolrich, M. W. & Smith, S. M. FSL. NeuroImage 62, 782–790 (2011).

84. Benjamini, Y. & Hochberg, Y. Controlling the False Discovery Rate: A Practical and Powerful Approach to Multiple Testing. J. R. Stat. Soc. Ser. B Methodol. 57, 289–300 (1995).

85. R: The R Project for Statistical Computing. https://www.r-project.org/.

86. Wood, S. mgcv: Mixed GAM Computation Vehicle with Automatic Smoothness Estimation. (2025).

87. Sedlacik, J. et al. Reversible, irreversible and effective transverse relaxation rates in normal aging brain at 3T. NeuroImage 84, 1032–1041 (2014).

88. Shaw, M., Rights, J. D., Sterba, S. S. & Flake, J. K. r2mlm: An R package calculating R-squared measures for multilevel models. Behav. Res. Methods 55, 1942–1964 (2023).

89. Hagler, D. J. et al. Image processing and analysis methods for the Adolescent Brain Cognitive Development Study. NeuroImage 202, 116091 (2019).

90. Feczko, E., Earl, E., Perrone, A. & Fair, D. ABCD-BIDS community collection (ABCC). Preprint at https://docs.abcdstudy.org/v/6_0_0/documentation/imaging/abcc_start_page.html (2020).

91. Feczko, E. et al. Adolescent Brain Cognitive Development (ABCD) Community MRI Collection and Utilities. 2021.07.09.451638 Preprint at https://doi.org/10.1101/2021.07.09.451638 (2021).

92. Glasser, M. F. et al. The minimal preprocessing pipelines for the Human Connectome Project. NeuroImage 80, 105–124 (2013).

93. Shafiei, G. et al. Reproducible Brain Charts: An open data resource for mapping brain development and its associations with mental health. BioRxiv Prepr. Serv. Biol. 2025.02.24.639850 (2025) doi:10.1101/2025.02.24.639850.

94. Fortin, J.-P. et al. Harmonization of multi-site diffusion tensor imaging data. NeuroImage 161, 149–170 (2017).

95. Johnson, W. E., Li, C. & Rabinovic, A. Adjusting batch effects in microarray expression data using empirical Bayes methods. Biostatistics 8, 118–127 (2007).

96. Fortin, J.-P. et al. Harmonization of cortical thickness measurements across scanners and sites. NeuroImage 167, 104–120 (2018).

97. Chen, A. A. et al. Mitigating site effects in covariance for machine learning in neuroimaging data. Hum. Brain Mapp. 43, 1179–1195 (2021).

98. Lenth, R. V. et al. emmeans: Estimated Marginal Means, aka Least-Squares Means. (2025).

99. Langkammer, C. et al. Quantitative MR Imaging of Brain Iron: A Postmortem Validation Study. Radiology 257, 455–462 (2010).

100. Bethlehem, R. a. I. et al. Brain charts for the human lifespan. Nature 604, 525–533 (2022).

101. Yu, M. et al. Statistical harmonization corrects site effects in functional connectivity measurements from multi-site fMRI data. Hum. Brain Mapp. 39, 4213–4227 (2018).

102. Péran, P. et al. Volume and iron content in basal ganglia and thalamus. Hum. Brain Mapp. 30, 2667–2675 (2009).

103. Ramos, P. et al. Iron levels in the human brain: A post-mortem study of anatomical region differences and age-related changes. J. Trace Elem. Med. Biol. 28, 13–17 (2014).

104. Larsen, B. et al. Maturation of the human striatal dopamine system revealed by PET and quantitative MRI. Nat. Commun. 11, 846 (2020).

105. Esteban, O. et al. FMRIPrep: a robust preprocessing pipeline for functional MRI. Nat. Methods 16, 111–116 (2019).

