## Supplemental Material for "A Scalable fMRI Estimate of Basal Ganglia Brain Tissue Iron for Use in Developmental and Translational Neuroscience"

**Supplementary Figure 1. Derived T2*w measures are highly similar across different processing configurations.** There was high similarity across T2*w measures produced by the 43 unique processing pipelines combinations. The absolute value Pearson’s correlations were computed between T2*w measures derived from each pair of processing pipelines across all participants, runs, and visits and all basal ganglia ROIs (618 total runs of T2*w data x 5 ROIs = 3090 T2*w measure datapoints for each pipeline). Pipelines differed in reference region used (corpus callosum (CC), ventricles (vent), whole brain, or a tSNR–informed whole-brain mask), in whether the mean or median was used to average signal across voxels within the reference region, in whether the mean or median was used to average voxelwise T2*w measure signal across volumes over time, in the normalization equations used to derive T2*w measure values (Equations 1–4), and in whether voxelwise scaling was applied or not (Equation 1 vs. Equation 2). Axis titles indicate the specific processing pipeline used to derive the T2*w measure, with components listed sequentially as: reference region (*CC*, *vent*, *wholebrain*, *tsnr_mask*); reference region averaging method (*mean*, *median*); time averaging method (*mean*, *median*); normalization equation with or without voxel scaling (Equation 1–4). The pipeline using a tsnr-informed whole-brain reference region had the highest similarity to all processing pipelines (mean *r* = .89).

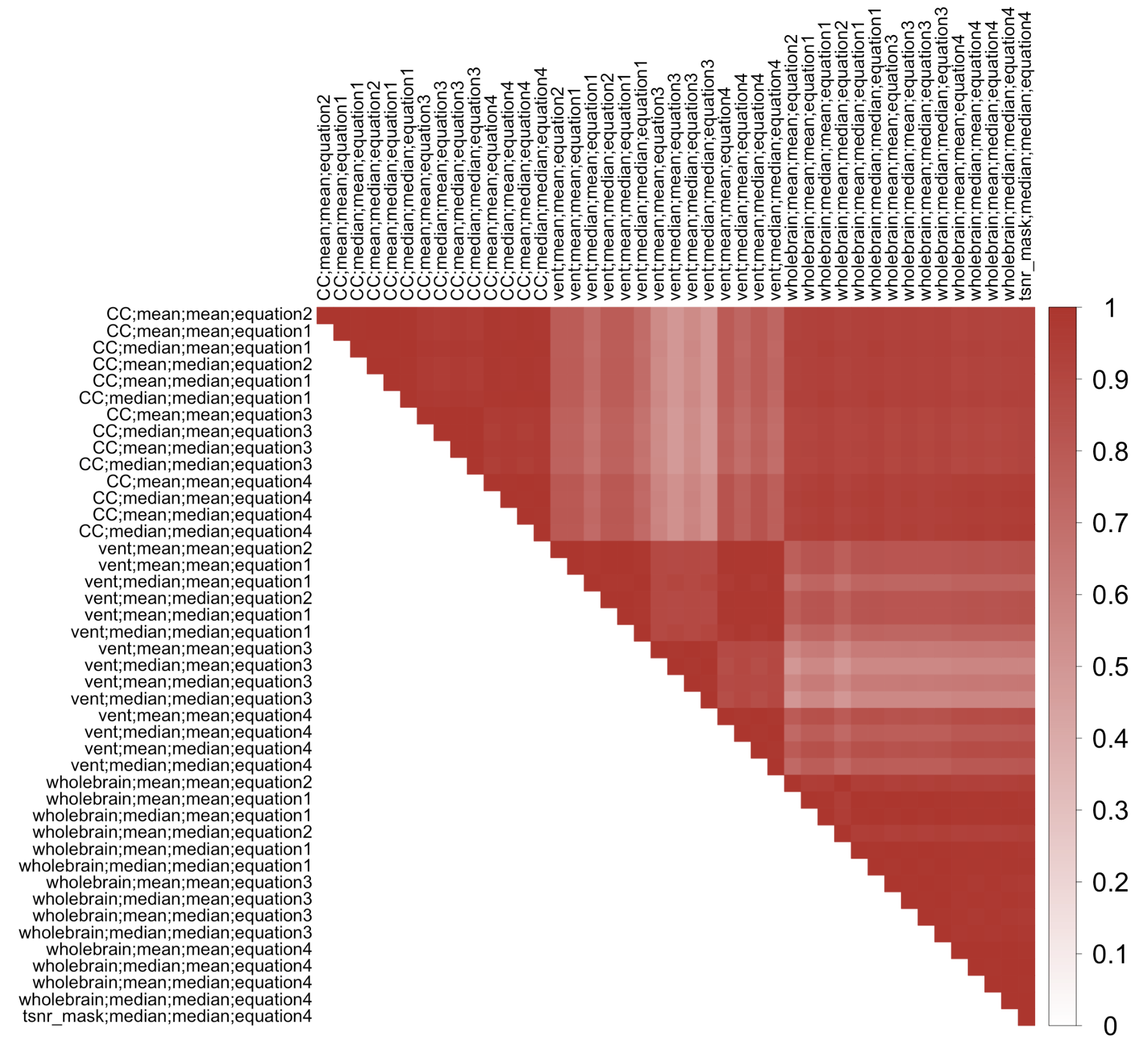

**Supplementary Figure 2. Deriving ΔR2* using a temporal signal-to-noise ratio restricted whole brain reference mask provides the best fit to standard quantitative relaxometry MRI measures.** Across all regions of the basal ganglia, the ΔR2* processing pipeline using a temporal signal-to-noise ratio (tSNR) -informed whole brain reference yielded a ΔR2* measure that explained the most variance in both A) R2* and B) R2', both gold-standard, relaxometry-based measures of brain iron.

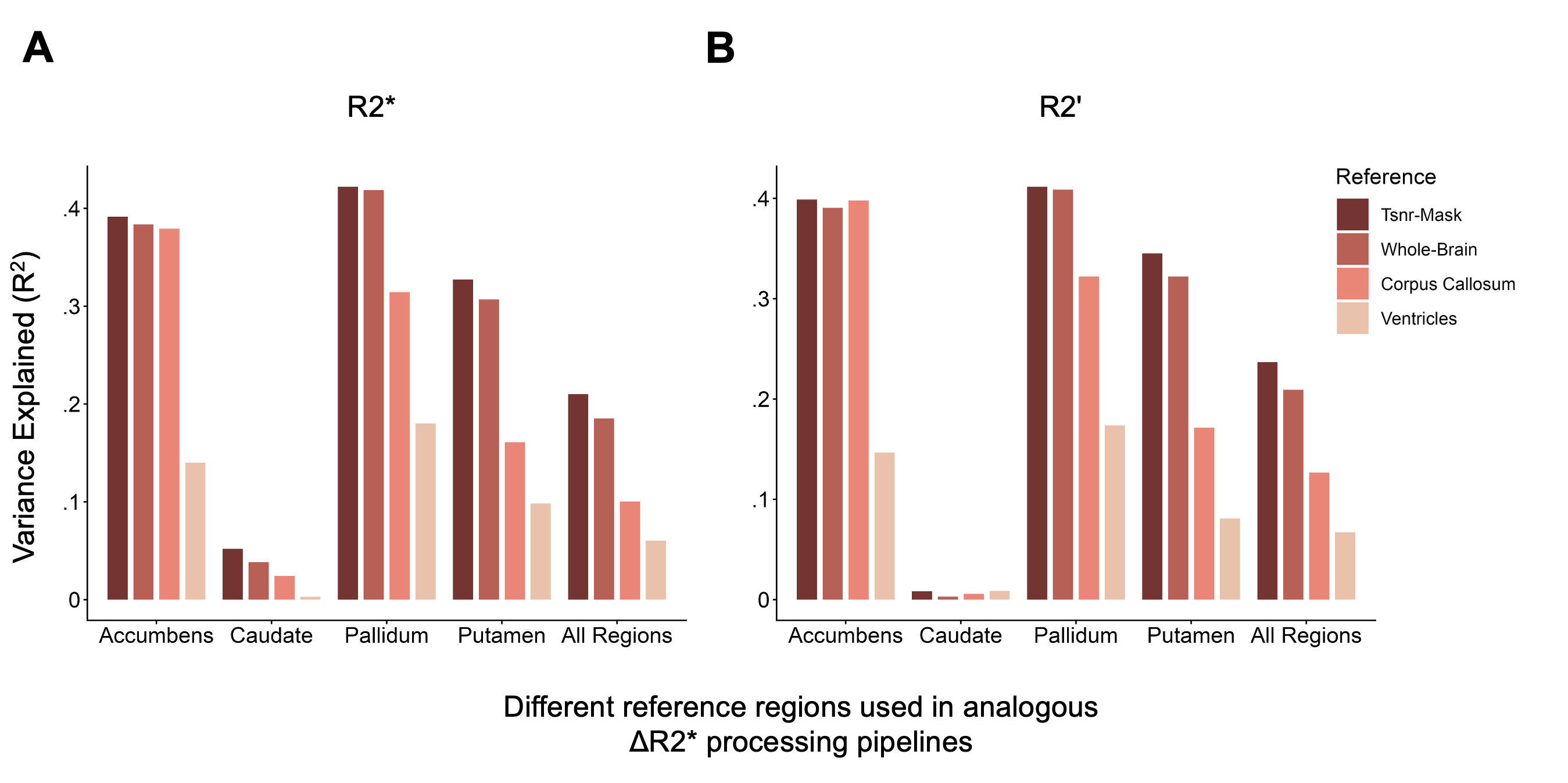

**Supplementary Figure 3. ΔR2* aligns with quantitative R2' relaxometry-based iron mapping across the basal ganglia.** To establish the validity of the ΔR2*, we compared ΔR2* maps derived from resting-state fMRI data to R2' relaxometry-based iron maps in a sample that contained both measures. A) The ΔR2* and R2' measures showed qualitatively similar spatial patterns with higher signal intensity (greater brightness) in iron-rich areas (e.g., the pallidum). B) ΔR2* and R2' measures show similar signal distributions and rank ordering across regions of the iron-rich basal ganglia, with the highest signal in the pallidum. C) ΔR2* is consistently associated with R2' across regions of the basal ganglia (*FDR* ≤ .001), after outlier removal on a per region level: accumbens n = 229; caudate n = 230; pallidum n = 230; putamen n = 230; all regions n = 229.

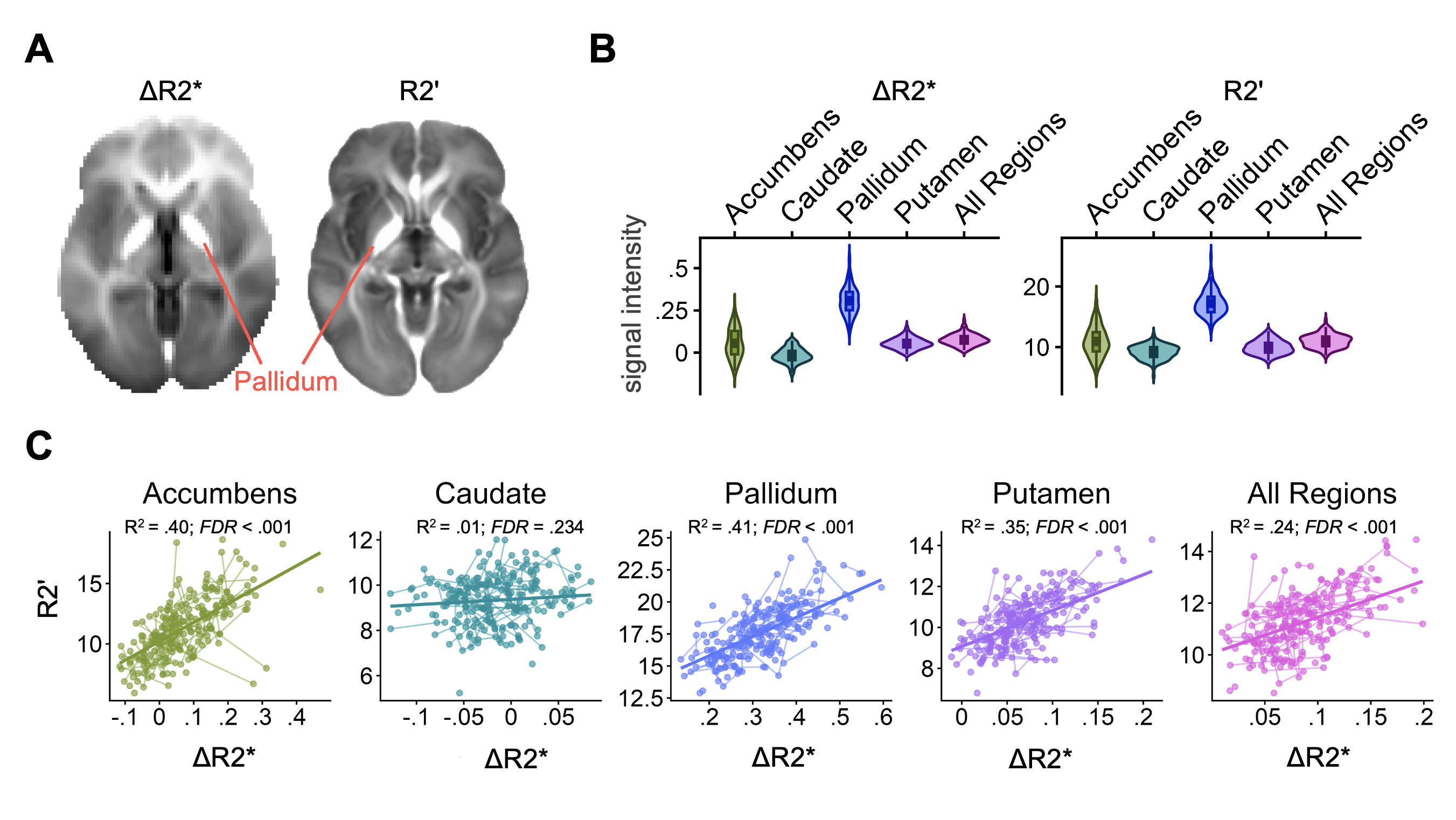

**Supplementary Table 1. Sample sizes (Ns) for analyses across basal ganglia ROIs.** Table values represent the total datapoints for analyses across basal ganglia regions following exclusions for data quality and (in Sample 1) outlier exclusion at the ROI-level.

|  |  |  | Accumbens | Caudate | Pallidum | Putamen | All Regions |
| --- | --- | --- | --- | --- | --- | --- | --- |
| **Validity (*R*^2^):** total sessions of data | | | | | |  |  |
|  |  | Against R2* | 229 | 232 | 231 | 229 | 229 |
|  |  | Against R2' | 229 | 230 | 230 | 230 | 229 |
| **Immediate test-retest reliability (*R*^2^):** total pairs of runs | | | | | |  |  |
|  |  | Visit 1 | 146 | 145 | 145 | 145 | 146 |
|  |  | Visit 2 | 98 | 99 | 98 | 99 | 99 |
|  |  | Visit 3 | 54 | 57 | 56 | 56 | 57 |
|  |  | All visits | 298 | 301 | 299 | 300 | 302 |
| **Longitudinal stability (*ICC*):** total within-run sessions of data  and total datapoints across all runs and sessions | | | | | | | |
|  |  | Run 1 | 264 | 267 | 263 | 266 | 266 |
|  |  | Run 2 | 259 | 261 | 259 | 260 | 262 |
|  |  | All runs | 523 | 528 | 522 | 526 | 528 |
| **All analyses in ABCD sample:** total participants (runs concatenated) | | | | | | | |
|  |  |  | 8366 | 8366 | 8366 | 8366 | 8366 |

**REFERENCES**

R Core Team, 2025: A Language and Environment for Statistical Computing. R Foundation for Statistical Computing, Vienna, Austria. <https://www.R-project.org/>

Wood, S.N. (2011). Fast stable restricted maximum likelihood and marginal likelihood estimation of semiparametric generalized linear models. *Journal of the Royal Statistical Society: Series B (Statistical Methodology)*, 73:3–36.

Simpson, G. L. (2018). Modelling Palaeoecological Time Series Using Generalised Additive Models. *Frontiers in Ecology and Evolution*, *6*, 396134. <https://doi.org/10.3389/fevo.2018.00149>

Simpson, G.L. & Singmann, H. (2025) gratia: Graceful ’ggplot’-Based Graphics and Other Functions for GAMs Fitted Using “mgcv” [Internet]. [cited 2025 Nov 24]. Available from: <https://cran.r-project.org/web/packages/gratia/index.html>

Benjamini, Y., & Hochberg, Y. (1995). Controlling the false discovery rate: A practical and powerful approach to multiple testing. *Journal of the Royal Statistical Society: Series B (Methodological)*, 57(1), 289–300.
